# Uncovering hidden genetic diversity: allelic resolution of insect and spider silk genes

**DOI:** 10.1101/2022.12.17.520845

**Authors:** Paul B. Frandsen, Scott Hotaling, Ashlyn Powell, Jacqueline Heckenhauer, Akito Y. Kawahara, Richard H. Baker, Cheryl Y. Hayashi, Blanca Ríos-Touma, Ralph Holzenthal, Steffen U. Pauls, Russell J. Stewart

## Abstract

Arthropod silk is vital to the evolutionary success of hundreds of thousands of species. The primary proteins in silks are often encoded by long, repetitive gene sequences. Until recently, sequencing and assembling these complex gene sequences has proven intractable given their repetitive structure. Here, using high-quality long-read sequencing, we show that there is extensive variation—both in terms of length and repeat motif order—between alleles of silk genes within individual arthropods. Further, this variation exists across two deep, independent origins of silk which diverged more than 500 million years ago—(1) the insect clade containing caddisflies and butterflies and (2) spiders. This remarkable convergence in previously overlooked patterns of allelic variation across multiple origins of silk suggests mechanisms for the generation and maintenance of structural protein-coding genes. Future genomic efforts to connect genotypes to phenotypes should account for such allelic variation.

## Intro

Silk is fundamental to the life histories of hundreds of thousands of arthropods (1). The genes that encode for silk proteins are often long and repetitive, and their variation is directly tied to silk phenotypes. Despite major headway in resolving difficult-to-assemble regions in *de novo* genome assembly [e.g., the human telomere-to-telomere consortium (2)], assemblies of biodiverse organisms vary widely in quality and contiguity (3, 4) and long repetitive regions pose a particularly difficult challenge (5). Arthropod silk genes are an example of a difficult assembly problem due to their repetitive internal region, which forms the semi-crystalline protein structure underlying the unique properties of silk (1). In the sister orders Lepidoptera (butterflies and moths) and Trichoptera (caddisflies), the gene that encodes the primary protein component of silk is *heavy chain fibroin (H-fibroin*), which originated in their common ancestor. However, structural silk protein-coding genes have independently evolved multiple times across the arthropod tree of life. For example, in spiders, there are multiple repetitive silk genes that encode a suite of proteins collectively known as spidroins (6).

The first full length *H-fibroin* sequence was published in 2000 from the model silkworm moth, *Bombyx mori*, through Sanger sequencing of a bacterial artificial chromosome (BAC) (7), and the first complete spidroin genes were published in 2007 using fosmid libraries (8). However, subsequent attempts using high throughput sequencing to resolve full-length *H-fibroin* and spidroin sequences largely failed due to the difficulty in resolving long repetitive regions with short-read sequencing (9). This lack of full-length fibroin and spidroin gene sequences has hindered the analysis of the variation present in these large (>20kbp), repetitive proteins, leaving a significant gap in our understanding of their evolution and function. Recently, advancements in long-read sequencing enabled recovery of these regions (10–13). However, only single consensus or haplotype sequences were recovered, leaving allelic variation completely hidden. New highly accurate long-read sequencing technologies, e.g., PacBio HiFi, can be assembled into haploid-resolved genomes, even for regions of the genome that were previously intractable to assemble (2, 5). Here we present the first examination of allelic structure in insect and spider silk genes and show that there is substantial diversity within individuals, demonstrating a wealth of genomic variation that was previously overlooked. Ultimately, characterizing and understanding this allelic variation is essential to understanding the molecular mechanisms that elaborated and shaped these highly modular structural proteins that are central to the success of hundreds of thousands of animals.

## Results and Discussion

We obtained high-quality reference genomes from single individuals of a butterfly (*Vanessa cardui*), three caddisflies from clades that span the breadth of the diversity of the order (*Hesperophylax magnus*, a case-making caddisfly; *Atopsyche davidsoni*, a cocoon-maker; and *Arctopsyche grandis*, a retreat-maker), and a spider (*Argiope argentata*). Two of these were newly generated and the primary assembles of the others were previously published (Table 1). These genomes represent some of the highest quality assemblies for their respective clades in terms of gene completeness (i.e., BUSCO scores) and contiguity (e.g., contig N50; Table 1).

**Table 1.** Comparison of genome and gene sequences from organisms used in this study. BUSCO scores reflect the insecta_odb10 dataset for the Trichoptera and Lepidoptera and the arachnida_odb10 dataset for the spider.

| Order | Species | Contig N50 (Mbp) | BUSCO % complete | Gene | Number of amino acids allele 1 | Number of amino acids allele 2 | Study |
| --- | --- | --- | --- | --- | --- | --- | --- |
| Trichoptera | *Arctopsyche grandis* | 6.5 | 98.2 | *H-fibroin* | 6375 | 5696 | Present study |
| Trichoptera | *Atopsyche davidsoni* | 14.1 | 98.8 | *H-fibroin* | 7878 | 6992 | Ríos-Touma et al. 2021 |
| Trichoptera | *Hesperophylax magnus* | 11.2 | 95.6 | *H-fibroin* | 8624 | 6728 | Hotaling et al. 2022 |
| Lepidoptera | *Vanessa cardui* | 7.0 | 98.8 | *H-fibroin* | 5675 | 4326 | Lohse et al. 2021 |
| Araneae | *Argiope argentata* | 32.3 | 98.4 | *MaSp2* | 4534 | 4183 | Present study |
| Araneae | *Argiope argentata* | 32.3 | 98.4 | *AgSp2* | 5465 | 5313 | Present study |

We recovered full-length, fully resolved sequences for both alleles of *H-fibroin* across all three caddisflies, unveiling substantial, and previously hidden, heterozygosity within each individual (Fig. 1). The variation between alleles can largely be ascribed to indels resulting in allele sequences with considerable differences in length (Table 1). Because the origin of *H-fibroin* can be traced back to the common, silk-spinning ancestor of Trichoptera and Lepidoptera more than 290 million years ago, we also investigated the *H-fibroin* sequence in the silk-spinning butterfly *V. cardui* to determine whether such patterns of heterozygosity were consistent across the evolutionary history of the gene. As with the caddisflies, we recovered two distinct *H-fibroin* alleles. Across all samples of Trichoptera and Lepidoptera, the structure of the *H-fibroin* was conspicuously conserved. For example, each *H-fibroin* sequence included conserved termini with a repetitive internal region consisting of modular repetitive units. Each gene was structured by a short initial exon followed by an intron and a long terminal exon that contained the entire repetitive region in a single open reading frame. Within each individual, allelic variation was dramatic, resulting from apparent deletions and insertions of repetitive modules (Fig. 1). However, despite the similarity in *H-fibroin* gene structure, the sequences and number of repeats in the internal regions varied widely across species and orders.

**Figure 1.**
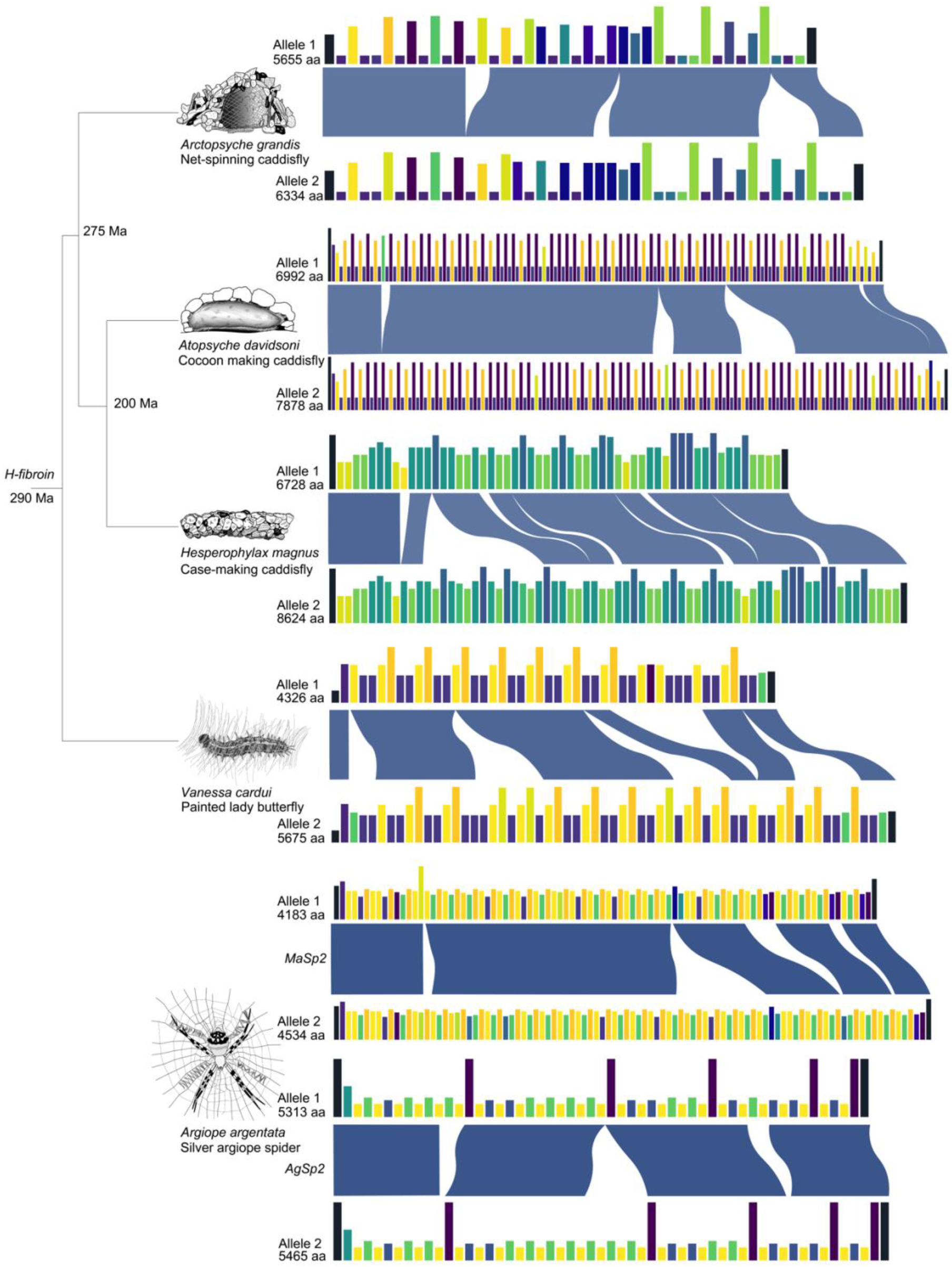
Allelic variation in silk genes of caddisflies, a butterfly, and a spider. Illustrations depict variation in silk use across these distantly related organisms. Allelic variation is shown for each gene. Bars of the same color represent the same repetitive motif within an individual, but not among individuals, and blue ribbons represent aligned regions highlighting insertions and deletions between alleles. The height of the bars indicates proportional unit length within each gene.

While the level of variation that we observed between alleles of *H-fibroin* within individual butterflies and caddisflies was striking, it is perhaps unsurprising that a functionally important gene with a common evolutionary origin would share similar patterns of variation. To determine if our findings extended to silk with a different evolutionary origin, we analyzed two spidroin silk genes from the spider *Argiope argentata: major ampullate spidroin-2 (MaSp2*) and *aggregate spidroin-2 (AgSp2*). For both, we observed patterns of allelic variation that were strikingly convergent with *H-fibroin*, including widespread modular heterozygosity between alleles (Fig. 1).

In this study, we uncovered remarkable convergence in the structure and variation between alleles of silk protein-coding genes across deeply divergent arthropod orders, including two independent origins of silk. Historically, this allelic variation has been overlooked for two reasons: first, short-read and/or low quality long-read sequencing are not well-suited to resolve long, highly repetitive regions, especially when there is extensive allelic variation (Hotaling et al. 2022). Second, single nucleotide variant (SNV) analysis, the most commonly way to assess genomic variation, often does not capture the full extent of variation in these repetitive regions. Now documented, the variation we observed lends insight into the mechanisms driving silk gene evolution (Fig. 1). For example, the allelic variation of large indels is consistent with unequal crossing over and gene conversion, both of which could play a role in driving allelic variation and motif homogenization. As such, selection may tolerate, and possibly favor, the presence of diverse alleles that differ in the organization of repeat units while maintaining nearly identical repetitive motifs at the nucleotide level. Furthermore, length variation among alleles may be critical to the function of silk fibers and represent the product of selection. However, to fully uncover the evolutionary mechanisms underlying variation in silk genes, population-level sampling and biomechanical testing are needed. Ultimately, to effectively connect genotype to phenotype [e.g., (13)], we must account for the full suite of allelic variation that exists within structural silk protein-coding genes. Our findings echo a broader trend; advances in long-read sequencing are leading to the discovery of substantial, and previously unobserved, genomic variation across the tree of life (14).

## Materials and Methods

We sequenced and assembled four of the five species represented here, two of which were available previously (Table 1). The other, *Vanessa cardui*, was publicly available (15). For each species, we extracted DNA from head and thorax (silk glands in the case of *Argiope argentata*) of a single individual using a Qiagen Genomic-tip extraction kit. After visualization of high molecular weight DNA using pulse-field gel electrophoresis, we sheared the DNA to 15 kbp fragments with the Diagenode Megaruptor. We generated PacBio HiFi libraries for each DNA extraction using the SMRTbell Express Template Prep Kit 2.0. Depending on genome size, each library was run on 30-hour 8M PacBio SMRT cells on the Sequel II instrument at the Brigham Young University DNA Sequencing Center.

We computed HiFi reads from the raw data (reads with quality above Q20) using PacBio SMRTlink software. We then assembled the raw reads into assemblies using HiCanu and Hifiasm v.0.13-r307 (16, 17). To extract and analyze the H-fibroin sequences, we used BLAST (18) to search the assembled genomes using the N-terminus and C-terminus sequences from previously published data. We then verified that both BLAST hits were isolated to the same contig in the genome assembly. We then extracted the region encompassed between the BLAST hits along with 20 kbp of flanking sequence on each end. We generated a gene annotation of the sequence using Augustus v.3.3.3 (19). The first species that we sequenced was *Hesperophylax magnus*. To generate assemblies for both *H-fibroin* alleles, we mapped HiFi reads back to the gene sequence from the primary assembly. We then separated reads into two bins by counting the number of mismatches in each read. We then assembled each bin of reads separately using HiCanu v.1.9 (17). Both assemblies resulted in full-length fully resolved *H-fibroin* sequences. We then verified that the allelic sequences matched those in the primary and alternate assemblies generated by Hifiasm (16). Both assemblers yielded identical sequences. For all other species, we simply extracted the alleles from the primary and alternate assemblies generated by Hifiasm (16) as described above.

We then aligned the amino acid sequences of the genes with the MAFFT-linsi algorithm to identify regions of shared alignment (20). To visualize the differences in repeat structure among species we generated a custom visualization script that splits the gene into repetitive motifs and then generates a bar plot, in which each unique motif is assigned a unique color (Fig. 1). To compare the nucleotide identity across motif classes, we wrote an additional script to extract the corresponding nucleotide sequences and then aligned them with MAFFT (20).

